# Development of a PC12 cell-based assay for *in vitro* screening of catechol-*O*-methyltransferase inhibitors

**DOI:** 10.1101/694596

**Authors:** Gongliang Zhang, Ingrid P. Buchler, Michael DePasquale, Michael Wormald, Gangling Liao, Huijun Wei, James C. Barrow, Gregory V. Carr

## Abstract

The male rat adrenal pheochromocytoma cell-originated PC12 cell line can synthesize and release catecholamine neurotransmitters, and it has been widely used as a model system in cell biology and toxicology research. Catechol-*O*-methyltransferase (COMT) is involved in the inactivation of the catecholamine neurotransmitters, and it is particularly important for theregulation of dopamine. In this study, we explored the feasibility of using PC12 cells as an *in vitro* drug screening platform to compare the activity of multiple COMT inhibitors. Incubation of PC12 cells with tolcapone, a highly potent and selective COMT inhibitor, increased the concentrations of dopamine and its metabolite 3,4-dihydroxyphenylacetic acid (DOPAC) while reducing the metabolites 3-methoxytyramine (3-MT) and homovanillic acid (HVA) in the cell culture medium. LIBD-3, a novel, non-nitrocatechol COMT inhibitor produced similar effects compared to tolcapone. LIBD-4, a less potent inhibitor, exhibited the expected right-shift in functional inhibition in the assay. These results match the known *in vivo* effects of COMT inhibition in rodents. Together, these data support the continued use of PC12 cells as an *in vitro* screen that bridges cell-free enzyme assays and more costly *in vivo* assays.

---

Dopamine (DA) is a neuromodulator that regulates motor function, reward-motivated behavior, cognition, and sympathetic tone.*^1^* The enzyme catechol-*O*-methyltransferase (COMT) metabolizes synaptically released DA and regulates dopaminergic signaling in the brain, particularly in the cerebral cortex and other regions with a low density of dopamine transporters (DAT)*^2, 3^* COMT inhibitors have been used as adjunctive therapies for Parkinson’s disease due to their ability to prevent the metabolism of L-DOPA in the periphery.*^4, 5^* Moreover, COMT inhibition has been proposed as a mechanism for improving cognition by tuning dopaminergic modulation of cortical function.*^6^* Because currently available COMT inhibitors have either insufficient brain penetration or severe toxicity, there is a need for novel, brain-penetrant, and safe compounds to validate the mechanism as a cognitive enhancement strategy.*^7^*

Preclinical *in vitro* and *in vivo* assays have been developed for dopaminergic drug screening. Cell-free *in vitro* assays are high-throughput and can provide valuable information on the effects of compounds on specific targets, but these assays provide no information on cell permeability, downstream effects of target engagement, or potential toxicity.*^8^ In vivo* microdialysis and fast-scan cyclic voltammetry provide information on the state of dopaminergic neurotransmission in whole animal systems, but are time-consuming, labor-intensive, and expensive, which makes them impractical for the early stages of drug discovery.*^3, 9, 10^* Cell lines that synthesize, release, and metabolize DA can potentially fill the gap between *in vivo* measures and cell-free systems.

Dopamine is synthesized in many cell types, including neurons and cells in the adrenal medulla.*^11^* PC12 is an established rat adrenal pheochromaocytoma-derived cell line that synthesizes and releases DA.*^12^* PC12 cells maintain a differentiated neuroendocrine phenotype and have been widely used as a model system for studies on neurotrophin action, protein trafficking, secretory vesicle dynamics, and neurotransmitter synthesis and release.*^12–14^* PC12 cells contain tyrosine hydroxylase*^15^*, monoamine oxidase (MAO)*^15^*, catechol-*O*-methyltransferase (COMT)*^16^*, and DA receptors.*^17^* Previous studies have demonstrated that PC12 cells release DA, epinephrine, norepinephrine, and acetylcholine in a calcium-dependent manner.*^18, 19^* These data indicate that PC12 cells have all of the required components for studying synthesis, synaptic release, and metabolism of DA. Taken together, these features suggest that PC12 cells may serve as a model of neuronal function with several advantages over primary neurons within the context of drug discovery, including increased consistency across batches and ease of production.

COMT inhibitors have been used to delineate the mechanisms of action for cytotoxic compounds in PC12 cells, but to our knowledge, there has not been extensive investigation of the effects of COMT inhibitors on DA release and metabolism in this cell line.*^20, 21^* To address this gap in the literature, we have conducted a series of experiments to validate a medium-throughput PC12 cell-based model of COMT inhibitor function.

## Results and Discussion

Figure 1A illustrates some of the critical steps in the synthesis and metabolism of dopamine and includes the enzymes that catalyze the individual reactions.*^22^* COMT and MAO are two enzymes involved in multiple steps in the dopamine metabolism pathway. COMT inhibition leads to decreases in 3-methyoxytyramine (3-MT) and homovanillic acid (HVA) concentrations and an increase in 3,4-dihydroxyphenylacetic acid (DOPAC) concentrations because it is the primary enzyme responsible for conversion of DA to 3-MT and DOPAC to HVA. There is substantial *in vivo* evidence that these neurochemical changes do in fact result from COMT inhibition.*^10, 23^* Conversely, MAO inhibition should lead to increases in 3-MT concentration because it is responsible for the conversion of 3-MT to 3-methoxy-4-hydroxyphenylacetaldehyde (MHPA) and decreases in DOPAC due to a decrease in 3,4-dihydrophenylacetaldehyde (DHPA), the substrate converted to DOPAC by aldehyde dehydrogenase.

**Figure 1.**
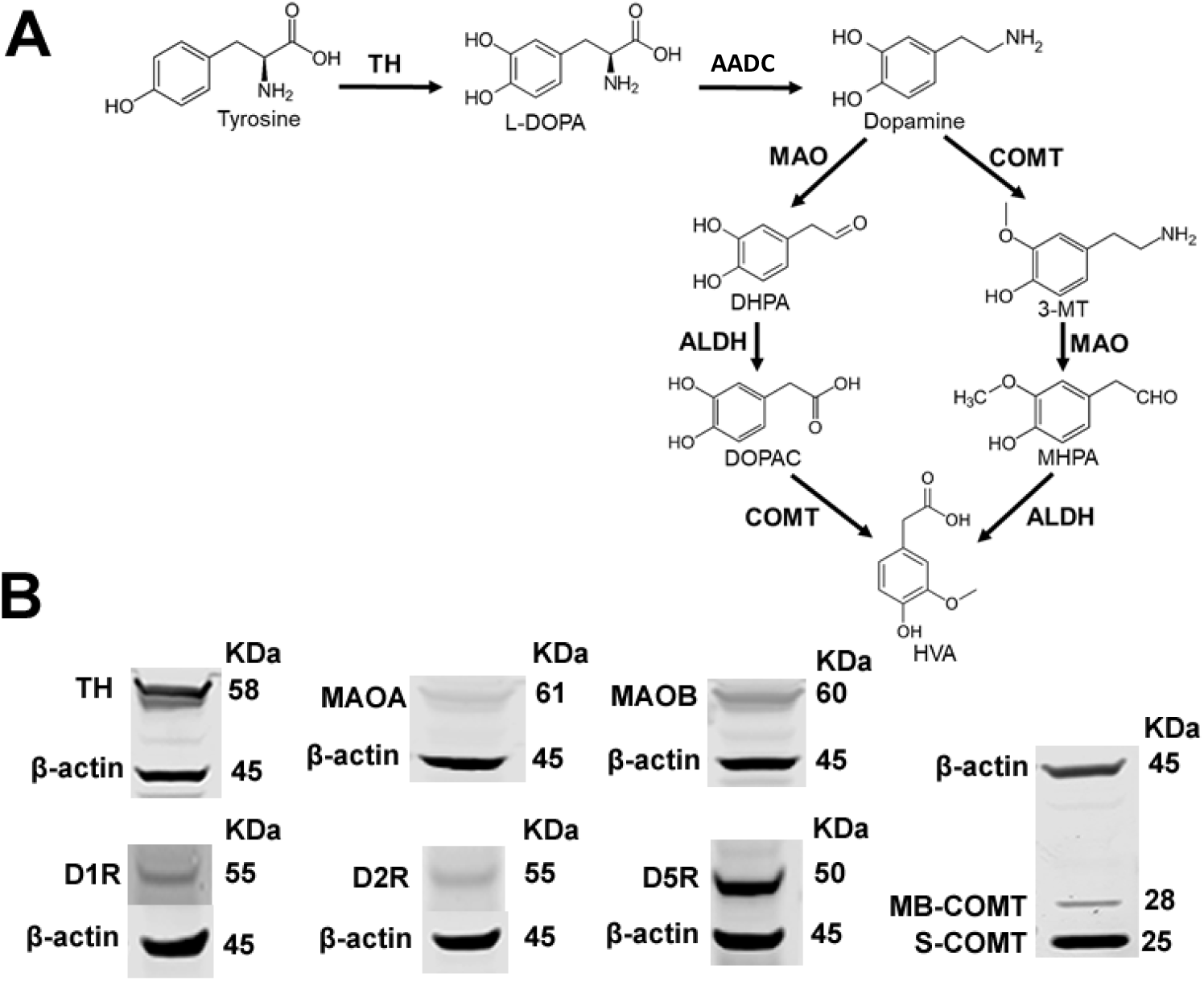
Proteins related to DA synthesis, release, function and metabolism are expressed by PC12 cells. (A) A schematic of DA metabolism. (B) Western blot studies revealed that PC12 cells presented TH, MAO-A, MAO-B, COMT, D1R, D2R and D5R. COMT blot shows membrane-bound COMT (MB-COMT) and soluble COMT (S-COMT). 3-MT, 3-methoxytyramine; ALDH, Aldehyde dehydrogenase. COMT, catechol-*O*-methyltransferase; AADC, aromatic-L-amino-acid decarboxylase; DHPA, 3,4-Dihydroxyphenylacetaldehyde; DOPAC, 3,4-dihydroxyphenylacetic acid; HVA, 3-methoxy-4-hydroxyphenylacetic acid or homovanillic acid; MAO, monoamine oxidase; MHPA, 3-Methoxy-4-hydroxyphenylacetaldehyde; TH, tyrosine hydroxylase.

Our first experiments were designed to confirm the expression of key regulators of dopaminergic function, including the COMT and MAO enzymes, in our PC12 cells. In agreement with previously published studies, we found expression of tyrosine hydroxylase, MAO, COMT, and D1, D2, and D5 dopamine receptors (Figure 1B). To determine if PC12 DA metabolite concentrations respond to COMT and MAO inhibition as expected, we utilized the COMT inhibitor tolcapone and the MAO inhibitor pargyline as positive controls. We also tested the effects of two of our novel COMT inhibitors, the highly potent LIBD-3 and the weak inhibitor LIBD-4 (Table 1).

**Table 1.** COMT inhibitor potency

| Compound | MB-COMT IC50 (nM) | S-COMT IC50 (nM) |
| --- | --- | --- |
| **Tolcapone** | 0.2 | 0.6 |
| **LIBD-3** | 1.5 | > 5000 |
| **LIBD-4** | 3767 | NT |

All COMT potency values are for recombinant enzyme (human)
NT = not tested

Because non-specific cytotoxic effects could modulate dopamine release and metabolism, we first tested the effects of our compounds on cell viability (Figure 2). Treatment with tolcapone, pargyline, and LIBD-4 up to 10 μM had no effect on viability. In contrast, 10 μM LIBD-3 significantly decreased viability, so all subsequent experiments with LIBD-3 only used concentrations up to 1 μM.

**Figure 2.**
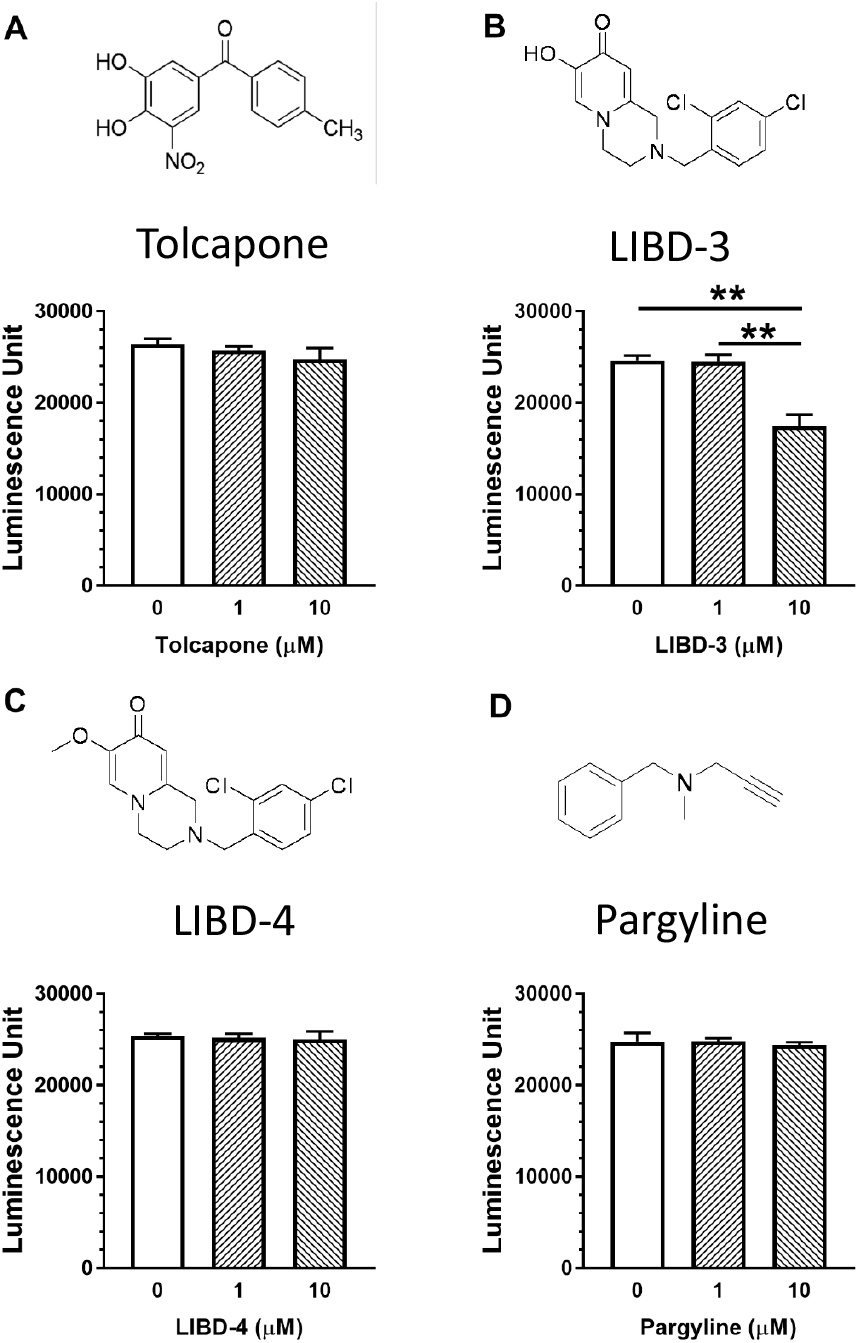
Cell viability assay. Test compounds at 0, 1 and 10 μM were incubated with PC12 cells for 24 hours. One-way ANOVAs did not any significant differences in the effects of tolcapone (A), LIBD-4 (C), pargyline (D) on cell viability. However LIBD-3 (B) at 10 μM, but not 1 μM, significantly reduced cell viability compared to vehicle treatment (*p* < 0.01, G). Data are expressed as mean ± SEM, n = 4 per group. ** *p* < 0.01.

We next measured the effects of 24-hour incubation of COMT and MAO inhibitors on the extracellular concentrations of DA and three metabolites, 3-MT, DOPAC, and HVA (Figure 3). We found that all three COMT inhibitors increased extracellular DA with LIBD-4, the least potent inhibitor increasing DA only at the highest drug concentration (10 μM). The MAO inhibitor pargyline did not alter DA concentrations at any dose tested (Figure 3A). The COMT inhibitor results differ from what has been reported for *in vivo* measurements which show that COMT inhibition alone does not appear to modify extracellular DA. This may be due to differences between a closed cell culture system and an open *in vivo* environment. DA is unable to diffuse out of the cell culture dish and PC12 cells do not endogenously express DAT*^24^*, so there is less capacity for DA reuptake and clearance from the extracellular space, potentially leading to accumulation. We measured the expected change in 3-MT for all compounds tested. The COMT inhibitors significantly decreased the 3-MT concentration while the MAO inhibitor pargyline increased it (Figure 3B). We also found the expected changes in DOPAC and HVA where the COMT inhibitors increased DOPAC and decreased HVA, while pargyline decreased both DOPAC and HVA. LIBD-4 produced effects at 10 μM, the only concentration above the IC_50_ of the compound, indicating that the PC12 model appears to give an accurate readout of target engagement (Figure 3C-D). These results demonstrate that COMT inhibitors produce changes in DA metabolite concentrations in PC12 cell cultures that are similar to those measured *in vivo*.

**Figure 3.**
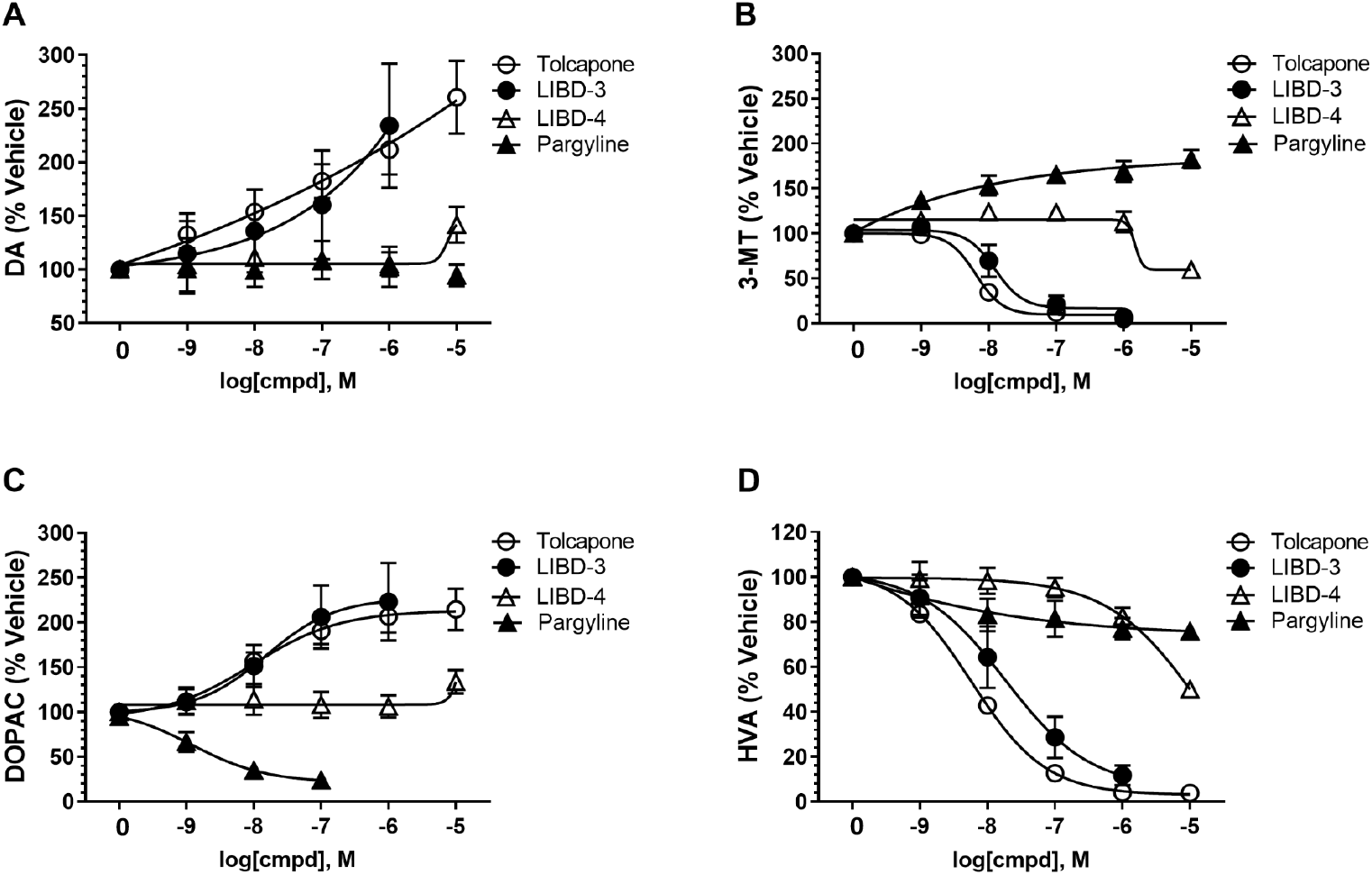
Effects of COMT and MAO inhibition on extracellular DA and metabolite concentrations/. PC12 cells were incubated with test compounds at concentrations of 0, 10^-9^, 10^-8^, 10^-7^, 10^-6^, and 10^-5^ M for 24 hours before collecting culture media. The extracellular concentrations of DA (A), 3-MT (B), DOPAC (C) and HVA (D) ions present in the culture medium in response to drug treatment are represented as a % of vehicle control. Tolcapone and LIBD-3 caused increases in DA and DOPAC levels and a decrease in 3-MT and HVA levels. LIBD-4 produced similar effects only at the 10^-5^ M concentration. Pargyline reduced DOPAC and increased 3-MT concentrations. Data are expressed as mean ± SD, n = 3 per group.

Microdialysis studies have shown that COMT inhibition augments extracellular DA concentrations in conjunction with high K^+^-induced depolarization.*^23^* We next tested whether a similar effect can be measured in PC12 cells. First, increasing the extracellular K^+^ concentration to 50 mM more than doubled the extracellular DA concentration compared to the levels present with 4.7 mM K^+^ (Figure 4A). Tolcapone and LIBD-3 (100 nM) both potentiated the extracellular DA concentration, but the changes compared to vehicle treatment did not reach our significance threshold (*p* = 0.0557 and 0.0518, respectively; Figure 4B). LIBD-4 and pargyline did not produce any change.

**Figure 4.**
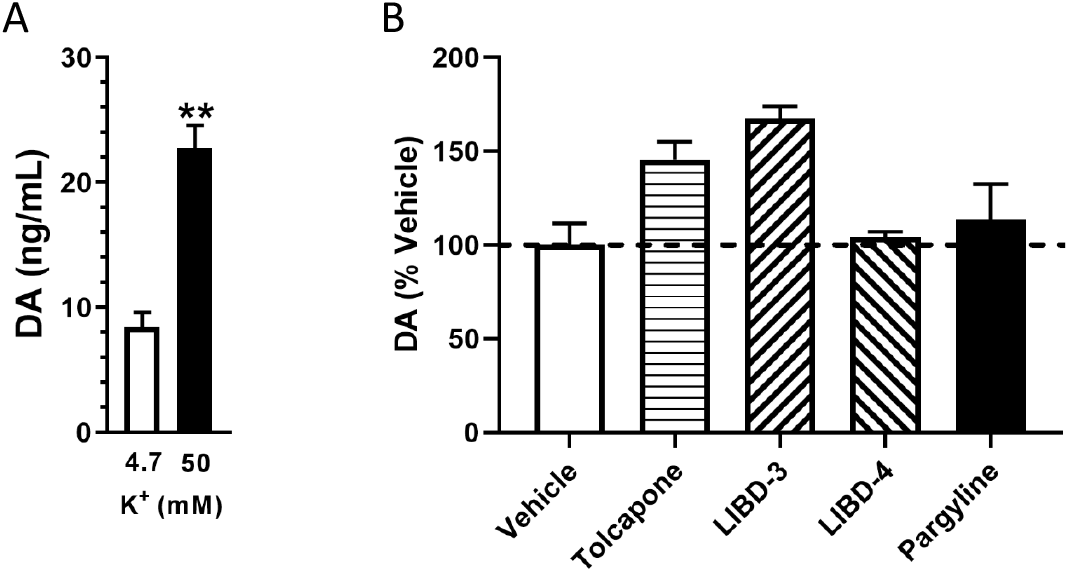
High K^+^ increases extracellular DA concentrations. (A) Increasing the extracellular K^+^ concentration leads to an increase in DA concentration in PC12 cells. (B) 100 nM Tolcapone and LIBD-3 appear to potentiate the effect of high K^+^, but the effects were not statistically significant (*p* = 0.0557 and 0.0518, respectively). LIBD-4 and pargyline at the same concentration had no effects. Data are expressed as mean ± SEM, n = 3 per group. ** *p* < 0.01.

The results described in these experiments demonstrate that PC12 cells can serve as a valuable *in vitro* model of COMT function and provide useful information on the relative activity of specific COMT inhibitors by measuring changes in DA metabolite concentrations. Similar effects of COMT inhibition on DA neurotransmission and metabolism have been demonstrated in previous *in vivo* experiments. However, the PC12 assays described in these studies provide similar information to more time-consuming *in vivo* studies without the use of animals and at a fraction of the cost. The higher throughput and greater overall efficiency of the PC12 assays allow for the collection of functional information on a greater number of COMT inhibitors at an earlier stage in the drug discovery process. This information can then be used to decipher the structure-activity relationship for novel chemical series more quickly and rank-order compounds for subsequent *in vivo* testing.

We compared the effects of two potent COMT inhibitors (tolcapone and LIBD-3) against those of a weak inhibitor (LIBD-4) and were able to measure significant differences, but with only three COMT inhibitors we were not able to investigate the level of correlation between the PC12 assay and *in vivo* assays.*^25^* We are currently completing a study with a large number of COMT inhibitors from multiple chemical series to determine if there is a good correlation in the magnitude of effects on DA metabolites between the PC12 cell model described here and an *in vivo* rat CSF sampling model.*^25, 26^* Additionally, the effects of COMT inhibition on high K^+^-stimulated DA release were not as clear as has been shown *in vivo^23, 27^*. This discrepancy may be due to the COMT inhibitor effects on baseline extracellular DA in PC12 cells or potential differences in clearance mechanisms.

Overall, we have found that PC12 cells represent a model of COMT function that can be used to screen COMT inhibitors and the resulting drug effects on DA metabolite concentrations mimic established *in vivo* models. Due to the high fidelity of the model system, PC12 cells can be used as a cost-effective bridge between cell-free enzyme assays and whole animal preparations.

## Methods

### Cell Culture and Media

PC12 cells were obtained from the American Type Culture Collection (Manassas, VA, USA; Product #: CRL-1721) and cultured in DMEM (Gibco/ThermoFisher Scientific; Product #: 31600-034) supplemented with 10% horse serum (Gibco/ThermoFisher Scientific; Product #: 16050-122), 5% fetal bovine serum (HyClone/ThermoFisher Scientific; Product #: SH30071.03), 1% penicillin streptomycin (Gibco/ThermoFisher Scientific; Product #: 15070-063) in a humidified incubator containing 5% CO_2_ at 37 °C. The maximal cell passage number was 15 to prevent change in cellular function.

#### COMT enzyme activity assay

COMT activity was measured using the MTase Glo methyltransferase assay (Promega, Madison, WI, USA) according to manufacturer’s instructions. Assays were carried out in Corning low volume 384-well white flat-bottom polystyrene NBS microplates with a final volume of 5 μL containing approximately 4 ng of human MB-COMT or 1 ng of human S-COMT as estimated by the Bradford Lowry method from the membrane homogenate. All reactions contained 20 μM high purity S-adenosyl methionine (SAM, CisBio, Bedford, MA, USA) in COMT assay buffer (50 mM Tris, 5–10 mM MgCl_2_, 2.5 mM DTT, pH 6.9). For MB-COMT, the catechol substrate was 7 μM norepinephrine (MilliporeSigma, St. Louis, MO,USA) and for S-COMT the substrate was 10 μM 7,8-dihydroxy-4-methylcoumarin (MilliporeSigma, St. Louis, MO, USA).

Reactions were performed in a 37 °C incubator for 1 h. The plate was removed from the incubator and allowed to cool to room temperature for 15 min. MTase reagent A (Promega) was first diluted 1:5 into RO water, and 1 μL was then added to the well. The plate was spun down, shaken, and allowed to incubate for 30 min at room temperature avoiding light. Then 5 μL of MTase reagent B (Promega) were added to all of the wells. The plate was spun down, shaken, and allowed to incubate for 30 min at room temperature avoiding light. Luminescence was detected with a Tecan Infinite M100 Pro plate reader.

#### Standard Curve

A standard curve was run on every plate. The amount of S-adenosyl homocysteine (SAH; CisBio) produced was determined using a standard curve and a linear back-calculation method. The standard curve comprised of varying concentrations of SAH from 500 nM down to 0 nM while maintaining a final SAM/SAH concentration of 50 μM. To correct for background levels present in the enzymatic lysate (MB-COMT), enzyme at assay concentration was added to the standard curve as well.

#### Determination of Inhibition

Percentage inhibition was calculated by using 10 μM tolcapone as the 100% inhibition value and the DMSO control as the 0% inhibition value. The dose response curves were constrained at 0% inhibition while keeping the percentage inhibition of the highest compound concentration floating. IC_50_ was determined by nonlinear regressions and curve fitting using a four-parameter fit with a variable slope in the Dotmatics studies program (Dotmatics, Bishops Stortford, UK). Potency data presented is an average of three separate experiments in which each data point was run in triplicate and reported as the IC50.

#### Cell Viability Assay

Drug toxicity was examined with CellTiter-Glo Luminescent Cell Viability Assay (Promega, G7572). In brief, PC12 cells were plated at a density of 5 × 10^4^ cells/100 μL/well in white opaque 96-well plates. Equal amounts of reagent was added into each well, followed by mixture for 2 min on an orbital shaker to induce cell lysis. The plate was equilibrated at room temperature for 30 min to stabilize luminescent signal before reading with a Synergy H1 Microplate Reader (BioTek, Winooski, VT).

### Western Blotting

The PC12 cell pellet was sonicated for 5 s in Pierce RIPA buffer (Thermo Scientific, 89900) containing 1% Halt protease & phosphatase inhibitor cocktail (Thermo Scientific, 1861281) on ice. Cell lysates were centrifuged at 10.000 × g for 15 min to pellet the cell debris. The supernatant was transferred to a new tube for protein quantification with a Pierce BCA Protein Assay Kit (Thermo Scientific, 23225). 40 μg of protein was separated by NuPAGE 4-12% Bis-Tris Protein Gels (Invitrogen, NP0335BOX) and transferred to a nitrocellulose membrane with an iBlot® Transfer Stack (ThermoFisher Scientific, IB301001). After blockade with Odyssey blocking buffer (LI-COR, 927-40000) for 1 h at room temperature, the membrane was incubated with the respective primary antibody (Table 2) at 4 °C overnight. After TBST washing, the membrane was incubated with the corresponding secondary antibody (IRDye 680LT and 800CW Infrared Dye, 1:15,000) for 1 h at room temperature. The western blot protein bands were captured by Odyssey CLX and analyzed by Image Studio software (V3.1, LI-COR Biosciences).

**Table 2.** Primary antibodies used in Western blot

| Antigen | Host | Dilution | Company |
| --- | --- | --- | --- |
| TH | Mouse, monoclonal | 1:10.000 | BD, 612300 |
| MAO-A | Rabbit, polyclonal | 1:100 | Santa Cruz, sc-20156 |
| MAO-B | Mouse, monoclonal | 1:500 | Santa Cruz, sc-515354 |
| COMT | Mouse, monoclonal | 1:5.000 | BD, 611970 |
| D1R | Mouse, monoclonal | 1:100 | Santa Cruz, sc-33660 |
| D2R | Mouse, monoclonal | 1:100 | Santa Cruz, sc-5303 |
| D5R | Rabbit, polyclonal | 1:1.000 | Proteintech, 20310-1-AP |
| β-actin | Rabbit, monoclonal | 1:1.000 | Cell Signaling, CST-13E15 |
| β-actin | Mouse, monoclonal | 1:5.000 | Abcam, ab8226 |

### Drug Treatment

PC12 cells were plated at a density of 5 × 10^4^ cells/100 μL/well in clear 96-well plate. 11 μL drug stock solution diluted in culture medium was added to corresponding well right after cell plating. After 24 hours incubation, the culture media was collected for analysis of DA and metabolites. In the high K^+^ (50 mM) induced DA release experiment, PC12 cells were cultured overnight in clear 96-well plates. The culture medium was removed and cells were washed with PBS one time before adding 70 μL K^+^ solution in PBS. The K^+^ solution was collected 15 min after incubation and centrifuged at 2.000 × g for 5 min to remove the cell pellet. All samples were stored at −80 °C before analysis. The formula of the high K+ solution consisted of (mM): NaCl 115, KCl 50, KH_2_PO_4_ 1.2, CaCl_2_ 2.5, MgSO_4_ 1.2, Glucose 11, and HEPES-Tris 15. Regular K+ solution consisted of (mM): NaCl 140, KCl 4.7, KH_2_PO_4_ 1.2, CaCl_2_ 2.5, MgSO_4_ 1.2, Glucose 11, and HEPES-Tris 15.

### Bioanalysis

#### Materials

DA, HVA, DOPAC, and 3-MT were obtained from MilliporeSigma (St. Louis, MO). DA-d3, HVA-d5, DOPAC-d5, and 3-MT-d4 were obtained from C/D/N Isotopes (Pointe-Claire, Quebec, CA).

#### Sample Preparation and Derivatization

Triplicate standards and QCs of dopamine, HVA, DOPAC, and 3-MT mixture were prepared in cell culture media. Triplicate 30 μL standards, QCs, and samples were transferred to the extraction plate. An equal volume of cold internal standard solution consisting of 500 ng/mL each of d0pamine-d3, HVA-d5, DOPAC-d5, and 3MT-d4 in 0.1% formic acid in acetonitrile (v/v) were added to each sample and the plate was mix for 5 minutes at 1250 rpm and centrifuged at 2650 x g for 20 minutes. 20 μL of supernatant was mixed with 10 μL of 100 mM sodium tetraborate in water and mixed for 1 minute. 10 μL of 2% benzoyl chloride in acetonitrile (v/v) was then added to each sample and the plate was mixed for 2 minutes at 1000 rpm. 10 μL of 1% formic acid in water was then added to each sample and mixed. 5 μL of each sample was injected for analysis.

#### LC-MS Analysis

The samples were analyzed using an Agilent 6540 QTOF with Jet Stream Electrospray Ionization Source (ESI) and Agilent 1290 UHPLC. Solvents were 10mM ammonium formate in water (A) and acetonitrile (B). Chromatographic separation was achieved over 7.5 minute using a Phenomenex Luna Omega 2.1 x 100 mm, 1.6 μm, C18 column with a binary gradient starting 21% B. The flow rate was 500 □ μL min-^1^. The autosampler was set at 20□°C. The LC gradient was as follows: 0 - 4□min, 21% B; 4 - 4.5□min, 21 - 40% B; 4.5 - 5.0 min, 40% B. 5.01 – 6.0 min, 95%B, 6.01 min, 21% B. The mass spec acquisition was performed using full scan MS from m/z 200 to 600 with the following source conditions: Drying and Sheath Gas temperatures at 350 °C and 400 °C, respectively; Both gas flows at 12 L/min; Nebulizer-45 psig, VCap, Nozzle and Fragmenter voltages at 3000 V, 600 V and 100 V, respectively.

Proton adducts of the triple benzoylated dopamine and dopamine-d3 (m/z 466.1649 and 469.1837, respectively) and the dual benzoylated 3-MT and 3-MT-d4 (m/z 376.1543 and 380.1794, respectively) as well as the ammonium adducts of the dual benzoylated DOPAC and DOPAC-d5 (m/z 394.1285 and 399.1599, respectively) and the single benzoylated HVA and HVA-d5 (m/z 304.1179 and 309.1493, respectively) were used for data analysis. Analyte peak areas were determined from the extracted ion chromatograms within a +/− 20 ppm window. Concentrations were calculated with linear regression analysis using Agilent Masshunter Quantitative Analysis Software (B.06.00 SP01).

#### Data analysis

All data represent three separate experiments with each data point from each experiment representing the average of two separate wells. Missing values on the curves are due to concentrations below the limits of quantitation (DA = 2.5 ng/mL; 3-MT = 2.5 ng/mL; DOPAC = 10 ng/mL; HVA = 20 ng/mL). One-way analysis of variance (ANOVA) was used to analyze the cell viability and high K^+^-stimulated DA release experiments with drug treatments. A paired t-test was used to analyze the low K^+^/high K^+^ comparison study. Significant main effects were further analyzed using Tukey’s *post hoc* test. All statistical tests were conducted using Prism 8.1.1 (GraphPad Software, Inc., San Diego, CA, USA). *p* < 0.05 was considered statistically significant.

## Abbreviations

3-MT: 3-methyoxytyramine
AADC: aromatic-L-amino-acid decarboxylase
COMT: catechol-*O*-methyltransferase
DA: dopamine
DAT: dopamine transporter
DHPA: 3,4-dihydrophenylacetaldehyde
DOPAC: 3,4-dihydroxyphenylacetic acid
HVA: homovanillic acid
MAO: monoamine oxidase
MHPA: 3-methoxy-4-hydroxyphenylacetaldehyde

## Author Information

### Author Contributions

GZ, MW, IPB, HW, JCB, and GVC designed the experiments. GZ, MD, MW, GL conducted the experiments. GZ, MD, IPB, HW, and GVC analyzed the data. All authors drafted and edited the manuscript.

